# A network of topographic numerosity maps in human association cortex

**DOI:** 10.1101/078931

**Authors:** Ben M Harvey, Serge O Dumoulin

## Abstract

Sensory and motor cortices each contain multiple topographic maps with the structure of sensory organs (such as the retina or cochlea) mapped onto the cortical surface. These sensory maps are hierarchically organized. For example, visual field maps contain neurons that represent increasing large parts of visual space with increasingly complex responses. Some visual neurons respond to stimuli with a particular numerosity, the number of objects in a set. We recently discovered a parietal topographic numerosity map where neural numerosity preferences progress gradually across the cortical surface, analogous to sensory maps. Following this analogy, we hypothesised that there may be multiple numerosity maps. Numerosity perception is implicated in many cognitive functions including foraging, multiple object tracking, dividing attention, decision making and mathematics. Here we use ultra-high-field (7T) fMRI and neural model-based analyses to reveal numerosity-selective neural populations organized into six widely separated topographic maps in each hemisphere. Although we describe subtle differences between these maps, their properties are very similar, unlike in sensory map hierarchies. These maps are found in areas implicated in object recognition, motion perception, attention control, decision-making and mathematics. Multiple numerosity maps may allow interactions with these many cognitive systems, suggesting a broad role for quantity processing in supporting many perceptual and cognitive functions.

---

Sensory and motor cortices each contain multiple topographic maps with the structure of sensory organs (such as the retina or cochlea) mapped onto the cortical surface. These sensory maps are hierarchically organized. For example, visual field maps contain neurons that represent increasing large parts of visual space with increasingly complex responses^1^. Some visual neurons respond to stimuli with a particular numerosity, the number of objects in a set. We recently discovered a parietal topographic numerosity map where neural numerosity preferences progress gradually across the cortical surface^2^, analogous to sensory maps. Following this analogy, we hypothesised that there may be multiple numerosity maps. Numerosity perception is implicated in many cognitive functions including foraging^3^, multiple object tracking^4^, dividing attention^5^, decision making^6^ and mathematics^7–9^. Here we use ultra-high-field (7T) fMRI and neural model-based analyses to reveal numerosity-selective neural populations organized into six widely separated topographic maps in each hemisphere. Although we describe subtle differences between these maps, their properties are very similar, unlike in sensory map hierarchies. These maps are found in areas implicated in object recognition, motion perception, attention control, decision-making and mathematics. Multiple numerosity maps may allow interactions with these many cognitive systems, suggesting a broad role for quantity processing in supporting many perceptual and cognitive functions.

Topographic maps have an orderly organization of neurons with similar functions. The close proximity of neurons with similar functions minimises local connection lengths to increase neural processing efficiency^10–12^. Furthermore, topographic maps allow simple one-to-one projections between maps. Finally, most neural processes are context-dependent. Topographic maps allow easy computations of context through comparisons between neighbouring neurons. Therefore, topographic organization has several benefits and gives a theoretical framework to explain why maps emerge in cognitive processing, as we recently demonstrated^2^ and extend here. Together with this numerosity map, we also demonstrated that another quantity, object size, is processed in a distinct object size map that largely overlaps with this parietal numerosity map, showing correlated numerosity and object size preferences^13^.

The parietal numerosity map we have described encompasses part of a network implicated in numerosity processing, extending into occipital, parietal and frontal areas^14–19^. The fine scale organization elsewhere in this numerosity network is unknown. We hypothesize that, like sensory maps, a hierarchy of several numerosity maps throughout human association cortices underlie this numerosity network. To investigate this hypothesis, we adapted our approach to reconstruct numerosity maps throughout the brain.

We displayed visual stimuli of changing numerosity while collecting ultra-high-field (7T) fMRI data covering the occipital, parietal, posterior-superior frontal and temporal lobes. We distinguished between responses to numerosity and co-varying stimulus features using several stimulus configurations^2, 16^. We summarized the fMRI responses using numerosity selective population receptive field (pRF) models with two parameters: preferred numerosity and tuning width. We consistently found six numerosity maps where these models explain responses very well (mean variance explained = 66%, corresponding to *p*=0.0022 (see Methods)) (Fig. 1a; Supplementary Fig. 1). These numerosity maps were often widely separated, but showed very similar patterns of responses. Each numerosity map contained very different responses separated by short distances (1-2 cm) (Fig. 1). Logarithmic Gaussian tuning functions explained slightly more response variance than linear Gaussian functions in all maps (Wilcoxon signed-rank test: NTO *p*<10^−20^, *z*=10.0, Δ=1%, n=1559; NPO *p*=10^−7^, *z*=5.2, Δ=0.1%, n=647; NPC1 *p*<10^−20^, *z*=29.0, Δ=1%, n=1675; NPC2 *p*<10^−20^, *z*=34.0, Δ=2%, n=1186; NPC3 *p*<10^−20^, *z*=16.6, Δ=1%, n=885; NF *p*<10^−20^, *z*=28.7, Δ=1%, n=1187, false discovery rate (FDR) corrected for multiple comparisons. Degrees of freedom (DF)=n-1), consistent with previous reports from some maps^2, 6, 13^.

**Figure 1:**
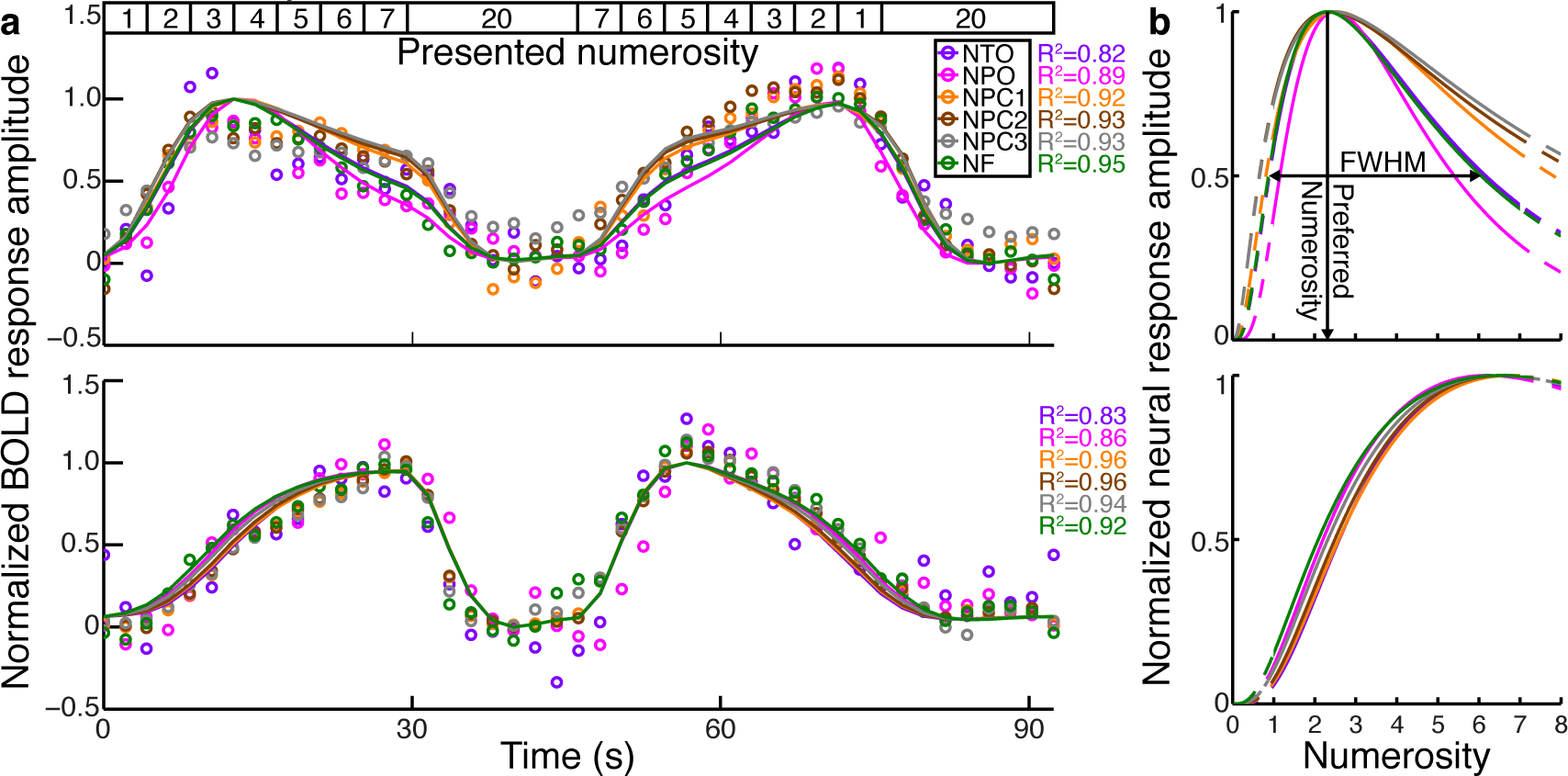
Similar responses to numerosity in several brain regions. (**a**) Varying stimulus numerosity (top inset) elicited different responses within each numerosity map (different colours). In the upper panel the largest response amplitude occurred after presentation of low numerosities, whereas in the lower panel they followed higher numerosities, considering the hemodynamic response delay. We captured these different responses using numerosity selective neural models, which captured high proportions of the variance (R^2^), in all time courses (R^2^>0.8). Between numerosity maps, responses and corresponding neural models were similar. Points represent response amplitudes averaged over all stimulus configurations; lines represent model predictions. (**b**) Representation of the neural model that best fits each time course. The model described a logarithmic Gaussian tuning function with two parameters: preferred numerosity and tuning width defined by the full-width-at-half-maximum (FWHM). Different model parameters explained the differences seen in the two panels in (a), capturing a similar proportions of response variance. Dashed lines are predicted neural responses beyond the tested stimulus range.

Projecting each recording site’s preferred numerosity onto the inflated cortical surface revealed six orderly topographic numerosity maps in each hemisphere (Fig. 2a, Supplementary Fig. 2). In each map, numerosity preferences changed gradually across the cortical surface, repeatably across subjects, scanning sessions and stimulus configurations (Supplementary Figs. 3 & 4). We named these numerosity maps after their anatomical locations, following extrastriate visual field map naming conventions^20^. We preceded their locations with ‘N’ for numerosity. Moving from posterior to anterior, the first numerosity map (NTO for ‘numerosity temporo-occipital’) lay at the lateral temporo-occipital junction, between the inferior temporal and lateral occipital sulci, posterior-superior to the preoccipital notch. NTO’s centre was at mean (SD) Montreal Neurological Institute (MNI) x,y,z coordinates 44(7), −75(1), − 4(3) in the right hemisphere and −42(3), −77(3), −3(8) in the left hemisphere. The second numerosity map (NPO) lay at the superior end of the parietooccipital sulcus (right 25(5), −82(4), 34(6), left −23(4), −80(5), 32(7)). The third, fourth and fifth numerosity maps (NPC1, NPC2 and NPC3) lay in and around the postcentral sulcus. NPC1 lay on the gyrus posterior to the superior postcentral sulcus (right 22(5), −61(7), 60(5), left −22(4), −59(11), 61(8)). NPC1’s location and orientation were very similar to our previous reports in the same subjects^2, 13^. NPC2 and NPC3 lay in the postcentral sulcus, superior and inferior (respectively) to its junction with the intraparietal sulcus (right 33(3), −40(4), 52(7), left −38(3), −43(8), 48(8) and right 45(10), −30(6), 40(4), left −48(6), −29(5), 34(6)). The sixth numerosity map (NF) lay at the junction of the precentral and superior frontal sulci (right 24(3), −11(5), 52(6), left −22(3), − 11(6), 50(8)). We found further numerosity selective areas in some cases, but not consistently between scanning sessions or subjects.

**Figure 2:**
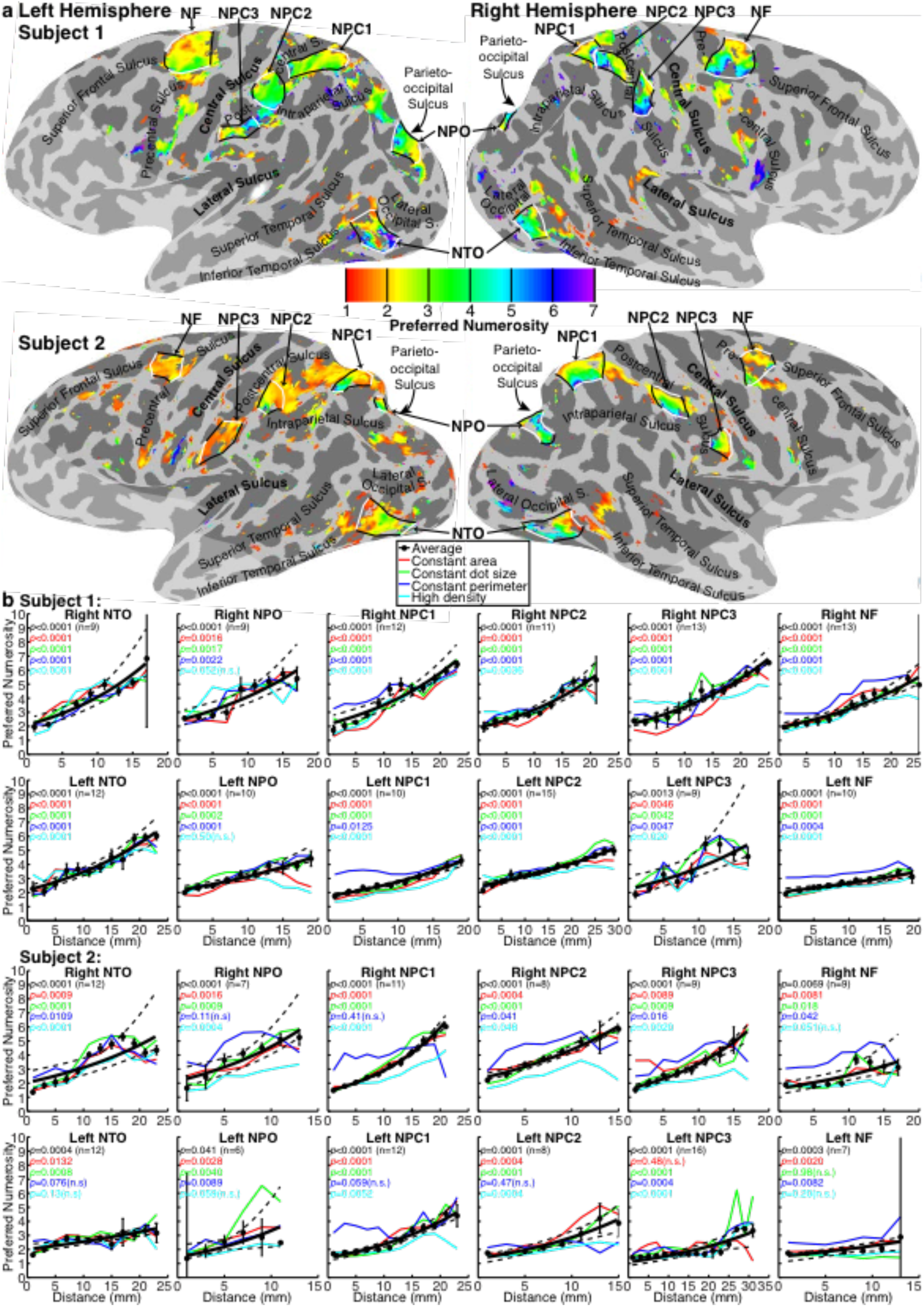
Numerosity map network. (**a**) Preferred numerosity in sites of significant numerosity-selective responses (variance explained > 30%, *p* < 0.0371 (FDR corrected for multiple comparisons)). There were several topographic numerosity maps, extended areas where preferred numerosity changed gradually across the cortical surface. Colours show each recording site’s preferred numerosity. White lines connect recording sites with the highest or lowest preferred numerosity present in each numerosity map. Black lines show borders of numerosity maps. Text labels show major sulci. Light shaded region was outside the fMRI recording volume. (**b**) Preferred numerosity of recording sites organized into bins by distance along each numerosity map’s cortical surface between the white lines in (a). In each numerosity map, preferred numerosity changed systematically and repeatably in each stimulus configuration. We fit distance bin mean preferred numerosities (black circles) with a logarithmic function (black line) with 95% confidence intervals to the fit (dashed black lines) determined by bootstrapping. Error bars show the standard error of the mean for each bin. Different stimulus configurations are represented as coloured lines joining the configuration-specific bin means. Coloured text gives probability of the observed change from permutation analysis, FDR corrected.

To quantify each numerosity map’s organization, we sorted each map’s recording sites by their distance to the map borders with the lowest and highest preferred numerosities present (the white lines in Fig. 2a & Supplementary Fig. 3). We then plotted preferred numerosity against cortical distance for each stimulus configuration and their average (Fig. 2b, Supplementary Figs. 4 & 5). In each numerosity map, preferred numerosity changed systematically and repeatably across the cortical surface in each stimulus configuration and scanning session, though less consistently in left hemisphere maps (Supplementary Fig. 5). We recorded each of 6 numerosity maps in 4 stimulus configurations in 5 subjects (n=120 measures per hemisphere). Permutation analysis revealed significant progressions of preferred numerosity with cortical distance in 107/120 right hemisphere measures and 98/120 left hemisphere measures (FDR corrected). A three-way ANOVA (n=240 numerosity map measures) revealed differences in the slope of the cortical numerosity progression between maps, stimulus configurations and hemispheres (map effect *p*=0.0001, F=5.7, DF=5; stimulus configuration effect *p*=4×10^−7^, F=11.6, DF=3; hemisphere effect *p*=0.015, F=6.0, DF=1). Subsequent multiple comparison tests^21, 22^ revealed that NF had less preferred numerosity progression than other maps, the constant perimeter stimulus configuration produced less preferred numerosity progression than other configurations, and left hemisphere maps had less preferred numerosity progression than right. Similarly, significant numerosity progressions were less frequent in NPO and NF than in other numerosity maps, in the constant perimeter and high density stimulus configurations, and in the left hemisphere (Supplementary Table 1).

Preferred numerosities within each numerosity map were well correlated between stimulus configurations recorded on different days (Supplementary Fig. 5), reflecting common topographical organization across stimulus configurations and repeated measures. However, numerosity preferences from the constant perimeter stimulus configuration were consistently less well correlated with preferences from other stimulus configurations^2, 13^.

The left and right hemispheres’ numerosity maps represented different numerosity ranges. Except in NTO, left hemisphere maps contained more recording sites with low preferred numerosities (below about three) than right hemisphere maps, while right hemisphere maps contained more high preferred numerosities (Figs. 2b & 3a). We use the upper quartile of preferred numerosities in each map to summarize this difference. A three-way ANOVA (n=60 numerosity maps) for effects of hemisphere, map, and subject first reveals that upper quartiles differ between hemispheres (*p*=2×10^−9^, F=54.0, DF=1). Subsequent multiple comparison tests revealed that right hemisphere upper quartiles were significantly higher than in left hemisphere, except in NTO (Fig. 3b). So right hemisphere numerosity maps typically had higher and broader distributions of preferred numerosity, though NTO had similar distributions across hemispheres.

**Figure 3:**
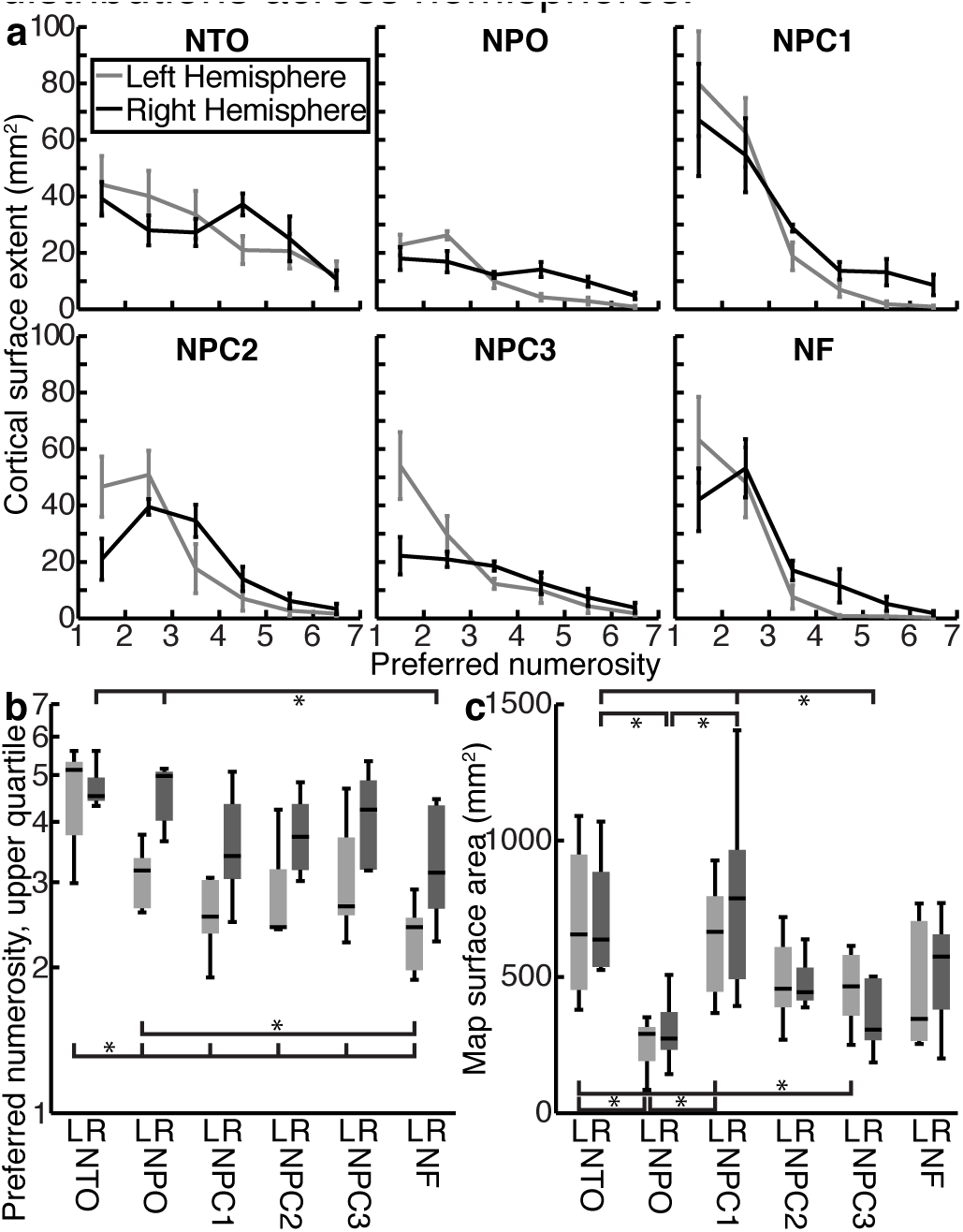
Differences in numerosity range and surface area between numerosity maps and hemispheres. (**a**) A larger proportion of each left hemisphere numerosity map had low preferred numerosities (below about three), while a larger proportion of each right hemisphere map had higher preferred numerosities. Error bars show the standard error of the mean. (**b**) Except in NTO, right hemisphere numerosity maps (dark grays) represented a higher range of numerosities than their paired left hemisphere maps (light grays), compared using the upper quartiles of the preferred numerosities present. There were also differences in the preferred numerosity distribution between numerosity maps and subjects within each hemisphere (see text). (**c**) Numerosity map surface areas differed between maps, but not between hemispheres. NTO and NPC1 were the largest numerosity maps, NPO was smallest, and other maps had similar intermediate surface areas. Horizontal lines are means, boxes are interquartile ranges of values from different subjects (not interquartile ranges of preferred numerosities), error bars are most extreme values. Brackets and stars show significant differences in subsequent multiple comparisons within each hemisphere: all brackets to the left of the star are significantly different from all brackets to the right of the star.

The same three-way ANOVA also revealed that upper quartiles of preferred numerosity distributions differed between subjects and between numerosity maps in the same hemisphere (map effect *p*=1×10^−8^, F=14.4, DF=5; subject effect *p*=3×10^−8^, F=15.6, DF=4) (Fig. 3b). Subsequent multiple comparison tests showed that left NTO had higher preferred numerosity distributions than other left hemisphere maps, which had similar distributions. More posterior right hemisphere maps (NTO and NPO) had significantly higher preferred numerosity distributions than anterior maps (NF). Right parietal maps (NPC1, NPC2 and NPC3) had intermediate preferred numerosity distributions that did not differ significantly from either posterior or anterior maps. So high numerosity preferences were primarily found in right posterior numerosity maps.

Numerosity map surface areas also differed (Fig. 3c). A three-way ANOVA (n=60 numerosity maps) revealed differences in map surface areas between maps and subjects, but not hemispheres (map effect *p*=4×10^−6^, F=9.0, DF=5; subject effect *p*=0.0011, F=5.4, DF=4). Subsequent multiple comparison tests within each hemisphere revealed that NTO and NPC1 were the largest numerosity maps, NPO and NPC3 were significantly smaller, and NPC2 and NF had intermediate surface areas. So differences between map surface areas did not suggest a progression from posterior to anterior.

In responses averaged across stimulus configurations, numerosity tuning widths consistently increased with preferred numerosity in each numerosity map (Fig. 4a, Supplementary Fig. 6). This increase was significant in 23/30 right hemisphere numerosity maps and 16/30 left hemisphere maps (FDR corrected).

**Figure 4:**
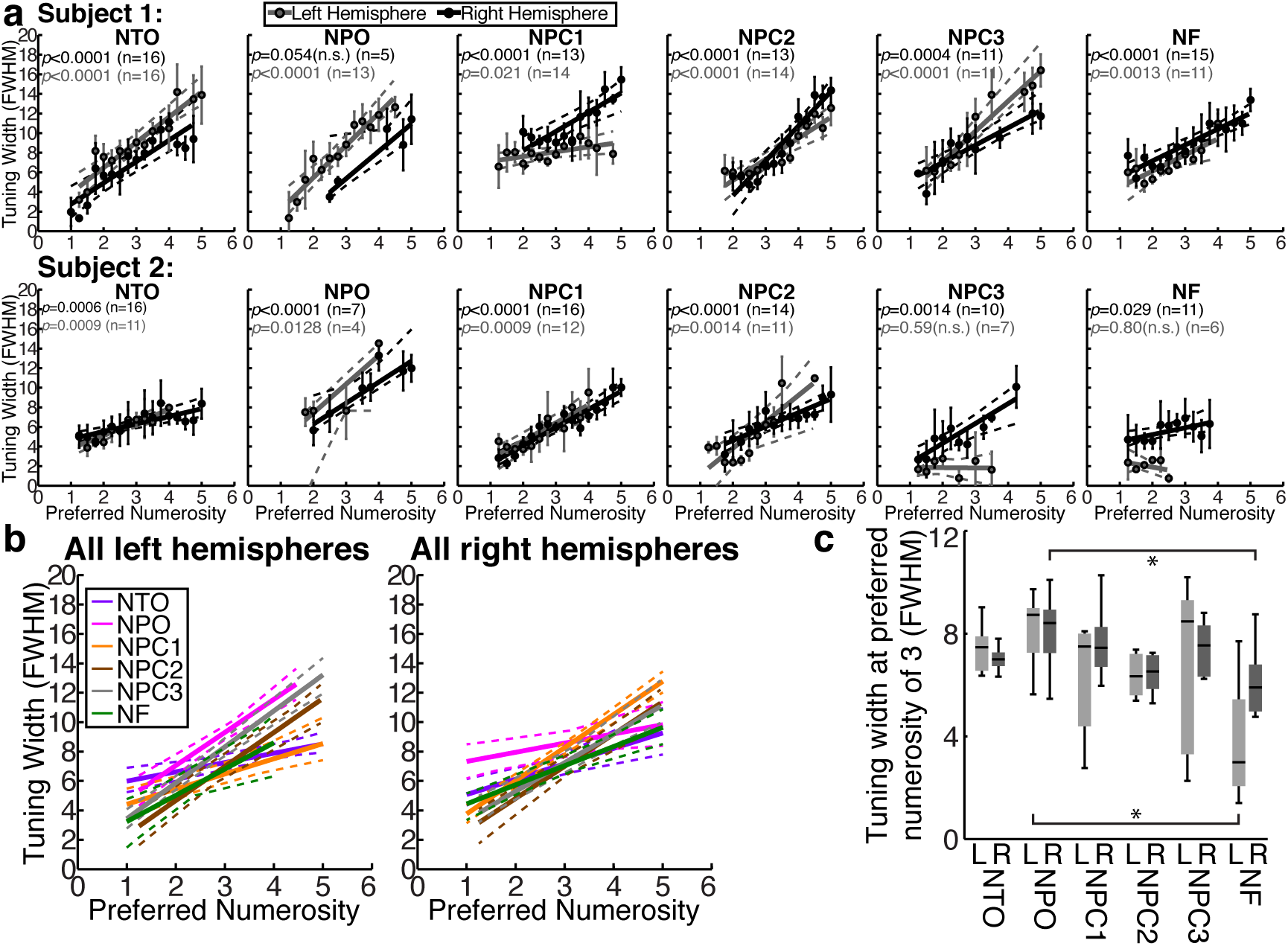
Numerosity tuning widths. (**a**) Progression of population tuning width with preferred numerosity in each numerosity map shown in Figure 2. We fit preferred numerosity bin mean tuning widths (black circles) with a linear function (black line) with 95% confidence intervals to the fit (dashed black lines) determined by bootstrapping. Error bars are standard errors. Text gives probability of the observed change from permutation analysis, FDR corrected. (**b**) Tuning widths grouped across subjects in each hemisphere. (**c**) Tuning widths differ slightly between maps in both hemispheres (see text). Brackets and stars show significant differences in subsequent multiple comparisons within each hemisphere: all brackets to the left of the star are significantly different from all brackets to the right of the star.

Maps in sensory cortex, and in particular visual cortex, show large systematic differences in tuning widths (i.e. receptive field sizes). Do numerosity tuning widths likewise differ between numerosity maps? To examine average tuning widths without biases arising from different preferred numerosity distributions, we quantified the fit tuning width progression at a preferred numerosity of three^23^. A three-way ANOVA (n=60 numerosity maps) reveals effects of map and subject, but not hemisphere, on tuning widths (map effect *p*=0.0015, F=4.6, DF=5; subject effect *p*=0.0011, F=5.4, DF=4) (Figs. 4b & 4c). Subsequent multiple comparison tests within each hemisphere reveal that NPO had the largest tuning widths, NF had significantly smaller tuning widths, and other maps had intermediate tuning widths that did not differ significantly from either NPO or NF. So there may be a slight tuning width decrease from posterior to anterior maps, but this is not as clear as the preferred numerosity distribution decrease.

Finally, we asked how numerosity map locations related to visual field map locations (Supplementary Figs. 7 & 8). All numerosity maps lay in or near visually responsive brain areas and partially overlap one or more extrastriate visual field maps. However, numerosity maps were not limited to the representation of the central visual field, where numerosity stimuli were presented. Furthermore, no numerosity map shared its borders with a visual field map. Finally relative locations of each numerosity map and its nearby visual field maps differed considerably between subjects and hemispheres. So we are confident that our numerosity maps do not reflect visual field map organization.

NTO overlaps with parts of visual field maps LO2, TO1 and/or TO2. Posterior superior NTO typically fell beyond of the parts of LO2 and TO1 covered by our visual field mapping stimulus. NPO overlapped with parts of IPS0 and/or IPS1. NPC1 overlapped with parts of IPS2, IPS3 and/or IPS4. NPC2 overlapped with parts of IPS4 and IPS5, often extending beyond IPS5 and beyond the eccentricity range we stimulated. NPC3 lay anterior and inferior to IPS5, beyond any previously described visual field map, but partially overlapping visual field position selective responses that suggest further undocumented inferior postcentral visual field map(s). Anterior NF typically partially overlapped with hFEF (the putative human frontal eye field map^24^).

This numerosity map network’s extent and overlap with other brain areas suggest human number and quantity processing may interact with several perceptual and cognitive functions. While processing in common areas implies interactions between neural systems^13, 25, 26^, different systems may be located together without interacting.

As our stimuli were presented visually, it is not surprising that responses were in visual areas. There are strong links between perception of numerosity and visual space or motor response location^27, 28^, likely mediated by working memory organization^29, 30^. So overlapping responses to numerosity (or its working memory footprint) and visual or motor location may underlie the cognitive spatial number line^2^. However, we find no clear relationship between numerosity and visual space preferences. Indeed, we find more high preferred numerosities in the right hemisphere, which primarily represents left visual and motor space, contrary to the usual association between high numbers and right visual or motor space^31^. So we find no obvious mechanism linking visual or motor space representations with numerosity representations.

The numerosity map network overlaps considerably with the fronto-parietal attention network of the intraparietal, postcentral and precentral sulci. Numerosity perception is more difficult with higher numerosities^6, 18^. Could tuned numerosity-selective responses reflect tuned responses to attentional load? This seems unlikely. First, our subjects performed no task with the displayed numerosities, and numerosity changed predictably. Second, attentional load should not differ in the constant perimeter stimulus configuration, but preferred numerosities differ here. Finally, tuned responses to attentional load have never been reported. So responses to numerosity do not straightforwardly reflect responses to attentional load.

Nevertheless, links between numerosity and attention seem likely. Display numerosity affects attentional object individuation capacity, suggesting numerosity representations guide attention’s spread between multiple objects^5^. Visual attention may use numerosity preferences to individuate objects and distribute attentional foci between them. Finally, stimuli attract attention, so numerosity-selective responses may reflect stimulus properties, or the distribution of attention the stimulus generates. Likewise, visual field position selective responses may reflect the position of the stimulus and/or the attention the stimulus attracts. Visual processing and attention affect each other, and attention may be an inherent component of stimulus-driven responses in higher visual processing.

Few recording sites had numerosity preferences above five. These numerosity preferences extend little beyond the subitizing range. While this low numerosity range is widely used to investigate numerosity selective responses in animals^6, 16, 17^, most human fMRI and behavioral studies use higher numerosities. The ‘approximate number system’ for higher numerosities depends less on attention^32^ and may not rely the system we characterize here^4, 33, 34^. The ability to decode numerosities beyond seven^14, 15^ suggests that responses to higher numerosities have spatial structure at fMRI resolutions. Alternatively, decoding of higher numerosities may depend on differential activation of sites with lower preferred numerosities.

NTO’s location in and near MT+ (visual field maps TO1 and TO2) and lateral occipital areas (LO1 and LO2) implicates numerosity in motion processing and object processing respectively. Interactions between numerosity selective and motion direction/speed selective populations may facilitate object individuation in multiple object tracking^4^. Likewise, the number of features within an object may help us perceive a face, or distinguish between rectangles and triangles.

Quantity processing may also guide decision making^6^ and support mathematical cognition^7–9^. The areas involved in these advanced and complex cognitive functions are less specifically localized, and seem to involve extensive networks that support several related functions. We do not map these, so cannot draw close links to areas supporting decision making and mathematical cognition. Nevertheless, the intraparietal sulcus (IPS) is often implicated in mathematical calculation^35, 36^ and both prefrontal and intraparietal areas in decision making^37, 38^.

Right hemisphere numerosity maps represent higher and broader numerosity ranges than left hemisphere maps, except NTO. This hemispheric difference supports previous reports of numerosity processing lateralization^2, 13, 39^, and extends this principle beyond NPC1 and the IPS. The proportion of high numerosity preferences present also decreases from posterior to anterior numerosity maps, though the functional significance of this trend is unclear. These differences result in small numerosity preference ranges in the left hemisphere’s anterior maps, with the upper quartile in left NF dropping as low as two. Therefore, numerosity preference progressions have very small effect sizes in some maps. However, recording sites with low numerosity preferences may contain information about higher numerosities: these sites may contain individual neurons with higher numerosity preferences, and neurons with low numerosity preferences respond to changes between higher numerosities.

Where numerosity preferences ranges and the slope of their cortical progression decrease, the slope and frequency of significant numerosity preference progressions also decreases, with only 60% of left NF measures showing a significant progression. Significant progressions are also less frequent in NPO, which has fewer recording sites from to quantify progressions. So differences in the slope and frequency of significant numerosity progressions (Supplementary Table 1) are linked to differences in preferred numerosity range and map surface area (Fig. 3).

The slope and frequency of significant numerosity preference progressions are lower in the constant perimeter stimulus configurations. Numerosity preferences measured with this configuration are also less well correlated with those from other configurations. We have previously shown that NPC1 voxels have object size preferences that are positively correlated with numerosity preferences^13^. The constant perimeter configuration shows small numerosities with very large object sizes and vice versa. Here, recording sites that prefer small numerosities and small object sizes may respond maximally to a larger numerosity because it has smaller objects. This reduces the numerosity preference progressions measured with stimulus configuration and the correlation to measures from other stimulus configurations. Finding this effect in outside NPC1 suggests other numerosity maps may likewise have object size selective responses.

Anterior visual field maps generally contain neurons with larger tuning widths (receptive field sizes) than posterior visual field maps. Other sensory processing hierarchies show similar progressions. Successive integration of visual information allows detection of more complex features by analysing successively larger areas of visual space. Numerosity tuning widths do not increase similarly, and indeed decrease slightly from posterior to anterior numerosity maps. Successive integration, if any, may therefore sharpen frontal numerosity representations. These are more closely linked to behavioural performance^40, 41^, so finer tuning widths here may improve behavioural performance. Alternatively, multiple numerosity maps may instead facilitate interactions with multiple perceptual and cognitive systems without successive integrations

Sensory and motor topographic maps are typically grouped in specific regions, allowing interactions over minimal distances. However, the numerosity maps are widely separated. This broad distribution may also facilitate interactions with several perceptual and cognitive systems.

Individual subjects differ in several numerosity map properties. Both map surface areas and the upper quartile of preferred numerosities differ between subjects. It is tempting to speculate that some individual difference in numerosity map properties might lead to some individual difference in numerical abilities. However, we remain sceptical of this link. Several behavioural measures (for example: subitizing range, accuracy, reaction time, or any measure of mathematical abilities) could be correlated with any numerosity map property (for example: numerosity map surface area, range of preferred numerosities, or tuning widths), in any of six maps in either hemisphere. Such analyses would also require far more subjects than we test here.

The positions of all numerosity maps are similar across subjects, as are the orientations of NPO and NPC1. However, the orientations of other maps vary. This variability resembles that of visual field maps. While early visual field maps are fairly consistently oriented, the frontal visual field maps hFEF and DLPFC show variable orientation between hemispheres and individuals^24^. However, both higher extrastriate visual field map and numerosity map orientations in each hemisphere are repeatable across independent scanning sessions, so we are confident of these orientations. We speculate that the increasing variability of anterior topographic map orientations may arise because these maps are not constrained by links to the orientations of neighbouring visual field maps and major neuronal pathways such as the optic radiation.

Some hemispheres contain multiple numerosity maps around a region where other hemispheres contain only one. This is clearest around NPC1, where further areas of numerosity selective responses were common. These may represent development of further numerosity maps is some subjects, particularly considering that the postcentral sulcus contains multiple numerosity maps in all subjects.

Macaque lateral prefrontal cortex (LPFC) and anterior inferior temporal cortex (AITC)^17^ recordings suggest numerosity selective responses in temporal and frontal areas that our scans do not cover. However, it is difficult to predict where homologous locations lie in macaques and in the greatly expanded human association cortices: macaque LPFC and AITC may be homologues of our NF and NTO maps respectively.

Macaque studies describe numerosity selective responses in the IPS, rather than the postcentral sulcus where we find three numerosity maps.

Macaques lack clearly distinguished postcentral and intraparietal sulci, so numerosity-selective responses in macaque IPS may be homologues of our NPC numerosity maps. FMRI numerosity mapping studies in macaques may clarify questions of homology, as they have in visual field mapping^42–44^.

Human fMRI studies consistently describe IPS activation during numerosity discriminations tasks^14, 18^, while our NPC numerosity maps lie in the postcentral sulcus. It is unclear whether previously described IPS locations correspond to NPC numerosity maps. Recent studies with careful response localization on the cortical surface show numerosity information at NPC map locations^15^, or at NPC map locations together with the IPS^14^.

However, IPS activation likely reflects responses to comparison tasks (which our subjects did not perform) rather than numerosity^45–47^. Furthermore, most studies use larger numerosities than the preferred numerosities we find.

Topographic organization is common to processing of sensory inputs, numerosity and other quantities. Numerosity maps form an extensive representation of quantity information throughout human association cortex. This includes areas involved in several cognitive and perceptual functions: visual motion processing, object recognition, attentional control, decision making and mathematics. As such, an extensive quantity processing system may form a major link between human perceptual systems and higher cognition.

## Methods

### Subjects

We present data from five human subjects (all male, aged 25-39 years). One was left handed. All were well educated, with good mathematical abilities. All had normal or corrected to normal visual acuity. All were trained with tasks requiring numerosity judgments before scanning. All gave written informed consent. These subjects were used in a previous study^13^, which included a small sub-set of this data. We have previously used the same number of subjects to characterize numerosity maps^13^. Our approach provides high statistical confidence in numerosity map locations and characteristics in each individual subject. We used multiple subjects to demonstrate reproducibility and generalization across subjects. We used four stimulus configurations on different days to demonstrate reproducibility over independent data collections. All experimental procedures were cleared by the ethics committee of University Medical Center Utrecht.

### Numerosity stimuli

Following protocols described in our previous studies^2, 13^, we presented visual stimuli by back-projection onto a 15.0×7.9 cm screen inside the MRI bore. Subjects viewed this through prisms and mirrors, and the subjects’ eyes were 41 cm from the display. Visible display resolution was 1024×538 pixels.

We generated stimuli the in Matlab using the PsychToolbox^48, 49^. A large diagonal cross of thin red lines crossed the entire display, facilitating accurate fixation at the cross intersection. Stimuli were groups of circles randomly positioned at each presentation so all circles fell entirely within 0.75° (radius) of fixation. To prevent perceptual grouping, individual circles were spread roughly homogeneously across this area (except in the high density condition described below).

We used various stimulus configurations^2^ to ensure low-level non-numerical stimulus features followed different time courses in different configurations. The first stimulus configuration (‘constant area’) kept summed circle surface areas constant across numerosities, ensuring equal luminance across numerosities. The second (‘constant object size’) kept individual circle size constant. The third (‘constant perimeter’) kept summed circle perimeters constant, ensuring equal edge extent across numerosities. The fourth (‘high density’) grouped the circles from the constant area figuration entirely within a 0.375° radius area that was randomly placed inside the stimulus area.

Patterns were black circles on a grey background. Patterns were presented briefly (300 ms) to make sequentially counting impossible. They were refreshed with a new random pattern every 700 ms, with 400 ms of gray background shown between pattern presentations. We presented each numerosity six times over 4200 ms (2 TRs), progressing slowly between numerosities. 10% of pattern presentations showed white circles instead of black. Subjects pressed a button when this happened, ensuring they were paying attention to the patterns. Subjects responded to 80-100% of white circle presentations in each scanning run. No numerosity judgments were required.

The numerosities one through seven were first presented in ascending order, followed by 16.8 seconds showing twenty circles, followed seven through one in descending order, followed by 16.8 seconds with twenty circles. We acquired 44 fMRI volumes during this cycle sequence, over 92.4 seconds, repeating this four times in each functional scanning run.

The long period of twenty circles allowed us to distinguish between very small and very large tuning widths^2, 50^. This is far outside the response range of neurons with small preferred numerosities, producing little neural response. Conversely, neurons responding to stimulus contrast energy should respond most strongly to numerous circles.

As in many fMRI experiments, these stimuli likely cause some adaptation to repeated presentation of the same numerosities^34, 51^. To minimize effects of adaptation on estimate numerosity preferences, we used a single model to summarize responses to both increasing and decreasing numerosity. This counterbalanced adaptation effects with stimuli that give both higher and lower responses preceding presentation of any numerosity, so reducing dependence on preceding stimuli.

### MRI acquisition and preprocessing

We acquired MRI data on a 7T Philips Achieve scanner. Acquisition and pre-processing protocols are described fully in our previous study^13^. Briefly, we acquired T1-weighted anatomical scans, automatically segmented these with Freesurfer, then manually edited labels to minimize segmentation errors using ITK-SNAP. This provided a highly accurate cortical surface model at the grey-white matter border to characterise cortical organization. We acquired T2*-weighted functional images using a 32 channel head coil at a resolution of 1.77×1.77×1.75 mm, with 41 interleaved slices of 128×128 voxels. The resulting field of view was 227×227×72 mm. TR was 2100 ms, TE was 25 ms, and flip angle was 70 degrees. We used a single shot gradient echo sequence with SENSE acceleration factor 3.0 and anterior-posterior encoding. Maximum gradient strength was 26 mT/m and maximum slew rate was 140 T/m/s. We used a 3rd-order image-based B0 shim of the functional scan’s field of view (in-house IDL software, v6.3, RSI, Boulder, CO, USA). This covered most of the brain, but omitted anterior frontal and temporal lobes, where 7T fMRI has low response amplitudes and large spatial distortions. Functional runs were each 176 time frames (369.6 seconds) in duration. The interval between runs was around one minute, the minimum possible. For each stimulus configuration, we acquired eight repeated runs in one session, with different configurations on different days.

We applied no spatial or temporal smoothing. We measured and corrected for head motion artefacts between and within functional scans. We then averaged functional data across each session’s runs, aligned it to anatomical scans and interpolated it into each subject’s anatomical segmentation space. We analysed responses from each configuration (session) separately and also averaged together.

### Code availability

We performed fMRI analysis using mrVista, which is freely available at (http://white.stanford.edu/software/). Custom code is added to this repository on publication.

### FMRI data analysis

We estimated numerosity response models from fMRI data and stimulus time courses as previously described^2, 13^, following a pRF modelling approach^50^. PRF models describe the aggregate tuning of neural populations within each grey matter fMRI recording site (n=159,136 recording sites across all subjects). Briefly, for each recording site and stimulus configuration, we used forward modelling to predict neural responses to the time course of displayed numerosities at each time point, for a set of candidate neural response models. We averaged all scanned stimulus cycles together, so each numerosity response model captured 44 fMRI response measurements (i.e. n=44). Each candidate neural response model described numerosity tuning using logarithmic Gaussian functions^2, 6, 13^ characterized by: (1) a preferred numerosity (mean of the Gaussian distribution) and (2) a tuning width (standard deviation of the Gaussian). Candidate neural response time courses reflected the overlap of the stimulus at each time point with this response model. We convolved these with a hemodynamic response function (HRF) to generate candidate fMRI time courses. For each fMRI recording point, we chose the tuning parameters whose associated fMRI time course most closely fit the recorded data, by minimizing the sum of squared errors (maximizing R^2^, variance explained) between the predicted and observed fMRI time series. Because HRF parameters differ between subjects, but differ little between brain areas or recording sessions^52^, we then estimated subject-specific HRF parameters across the whole acquired fMRI volume from all the data recorded from each subject, as described elsewhere^2, 23^, and re-fit response models using these HRF parameters.

We demonstrate that logarithmic Gaussian functions explain more response variance than linear Gaussian functions by also fitting pRF models where candidate neural response models were linear Gaussians^2, 6, 13^. We then compared the variance explained in all recording sites in each ROI (grouped across all subjects) using a Wilcoxon signed-rank test.

To convert these variance explained measures to probabilities of observing these model fits by chance, we using the same procedure to fit tuning models to recordings from 243,000 white matter recording points in the same scans, creating a null distribution^13^. We then determined the proportion of fits exceeding any variance explained. We used false discovery rate (FDR) correction for multiple comparision^53^, taking all grey matter recording sites in all subject’s scanning volumes into account.

We excluded recording sites where pRF models explained less than 30% of response variance (i.e. with a probability above 0.0371 of observing this goodness of fit by chance, FDR corrected) from further analysis.

Candidate preferred numerosities extended beyond the presented numerosity range, allowing model fit parameters beyond this range. This meant returned parameters within the stimulus range were reported accurately, not just the best fit of a limited set. We could not accurately estimate preferred numerosities outside of the stimulus range, so excluded any recording sites with preferred numerosities outside this range from further analysis.

### ROI definitions

Here we aimed to search for new numerosity maps throughout the cortical surface. However, our approach followed that in our previous demonstration of numerosity maps, which focused on the posterior parietal and dorsal occipital lobes^2^. Again, we started with the numerosity-selective response model from the average of all stimulus configurations. We rendered the variance explained of all recording points with preferred numerosities in the presented range onto the cortical surface (Supplementary Fig. 1). This highlighted six regions where numerosity selective response models consistently captured responses well. These regions were located similarly (with respect to major sulci) in all hemispheres, and formed the basis our regions of interest (ROIs). We then rendered the preferred numerosities of each recording site on the cortical surface around these regions (Supplementary Fig. 2). Across hemispheres and stimulus configurations, we consistently found topographic representations of preferred numerosity at these locations (6 maps x 2 hemispheres x 5 subjects = 60 maps, each measured repeatedly in 4 stimulus configurations on different days). In each map, we defined lines joining locations with equal preferred numerosity at the low and high ends of the preferred numerosity range seen in each numerosity map (the ‘ends’ of the map). The other map borders (the ‘sides’) followed the edges of topographic organization, where the goodness of model fits decreased.

### Conversion to MNI coordinates

Our analyses were in individual subject space. To describe map locations on an average brain, we converted these to Montreal Neurological Institute (MNI) x, y, and z coordinates. We first located each individual subject’s map centres on the cortical surface. We then transformed each subject’s anatomical MRI data, together with these map centre locations, into MNI average template space with MINC’s ‘mincresample’ tool (http://packages.bic.mni.mcgill.ca) using rigid alignment and linear scaling. We took the mean and standard deviation of the resulting MNI coordinates of each map across subjects.

### Analysis of changes across the ROI

We calculated the distance along the cortical surface from each point in each ROI to the nearest point on the lines of the lowest and highest preferred numerosities. The ratio between the distances to each line gave a normalized distance along the ROI in the primary direction of change of numerosity preferences. We multiplied this by the mean ROI length in this direction.

We binned the recording points within every 2 mm along each numerosity map, calculating the mean and standard error of their preferred numerosities in each stimulus configuration and their average. Bins were excluded if their cortical surface extent was smaller than one fMRI voxel or smaller than the point spread function of 7T fMRI^54^. Bins counts varied with the maps’ cortical extents, ranging from 6 to 18, and is given in each figure. We fit the best fitting logarithmic functions to bootstrapped samples of the bin means, because preferred numerosity progressions are fit better by logarithmic functions than straight lines^2, 13^. From these bootstrapped fits, we took the median slope and intercept as the best fitting numerosity progressions. We determined 95% confidence intervals by plotting all bootstrapped fit lines and finding the 2.5% and 97.5% percentiles of their values.

We determined the statistical significance of these slopes with a permutation analysis. We repeatedly (10,000 times) randomized which preferred numerosity was associated with each distance bin and fit the slopes of each permutation. We then determined how many of these permutations had equal or greater slopes than the observed data, the probability of observing this slope by chance. We FDR corrected this probability, taking into account probabilities from all stimulus configurations, maps, hemispheres and subjects.

Similarly, for each ROI we looked for changes in tuning width with preferred numerosity^2, 6, 13^. We binned the recording sites within every 0.25 increase in preferred numerosity, calculating the mean and standard error of their tuning width. Bins were excluded if their cortical surface extent was smaller than one fMRI voxel or smaller than the point spread function of 7T fMRI^54^. Few recording sites had preferred numerosities above 5, so we only used bins from 1 to 5 preferred numerosity. This formed a maximum of 17 preferred numerosity bins for each numerosity map, the number used in each map is given in each figure. We fit linear functions to bootstrapped samples of bin means and took the median slope and intercept as the best fitting tuning width progression. We used the permutation analysis described above to calculate the probability of observing this tuning width increase by chance.

### Analysis of differences between stimulus configurations and numerosity maps

We consistently found preferred numerosity progressions across the cortical surface in all stimulus configurations. However, numerosity preferences may vary between stimulus configurations, for example due to object size selectivity in the same recording sites^2, 13^. We grouped recording sites from the same numerosity map and hemisphere across all subjects. Recording sites counts differed between maps and hemispheres, and are given on Supplementary Figure 5. In each group of recording sites, we correlated the numerosity preferences measured in each stimulus configuration with each other configuration using Pearson’s correlation.

We tested whether the rate of preferred numerosity progression across the cortical surface was affected by stimulus configuration, numerosity map and hemisphere. For each subject, stimulus configuration, numerosity map and hemisphere, we took the slope of the best fitting logarithmic function to bins of preferred numerosity versus normalized cortical distance. We normalize cortical distance measures by total map length to avoid effects of map size differences between subjects and numerosity maps. We assessed differences between stimulus configurations, numerosity maps and hemispheres using a three-way analysis of variance (ANOVA) on these slopes (n=240). We revealed where these differences reach significance using subsequent multiple comparison tests^21, 22^.

We then tested whether the distribution of preferred numerosities within each numerosity map was affected by hemispheric lateralization, numerosity map identity and subject identity. For each numerosity map in each subject, we took each recording site’s preferred numerosity from responses averaged across all stimulus configurations. To summarize the range of numerosity preferences with a single number, we then determined the upper quartiles of preferred numerosities present in each numerosity map in each subject and hemisphere. We previously assessed interquartile range differences, but lower quartiles did not differ significantly between hemispheres or numerosity maps, so interquartile range differences primarily reflected upper quartile differences.

We then assessed differences in the distribution of preferred numerosities across hemispheres, subjects and maps using a three-way analysis of variance (ANOVA) on the upper quartiles of all maps (n=60). We revealed where these differences between hemispheres and maps reach significance using subsequent multiple comparison tests.

We similarly assessed numerosity map surface area differences by determining the area of each numerosity map on the folded cortical surface. Again, we assessed differences in the distribution of preferred numerosities across hemispheres, subjects and maps using a three-way ANOVA on these surface areas. Again, we used subsequent multiple comparison tests to reveal which numerosity maps differ significantly in surface area.

Finally, we assessed tuning width differences between numerosity maps. Numerosity tuning widths vary systematically with preferred numerosity, and preferred numerosity distributions differ between maps. To examine differences in average tuning width without biases arising from different preferred numerosity distributions, we determined the value of the fit tuning width progression for a preferred numerosity of three, an intermediate preferred numerosity that was present in almost all numerosity maps^23^. Again, we assessed tuning width differences across hemispheres, subjects and maps using a three-way ANOVA. Again, we used subsequent multiple comparison tests to reveal which numerosity maps differ significantly in tuning width.

### Visual field mapping

We acquired visual field mapping responses to examine the relationship between numerosity maps and visual field maps. The visual field mapping paradigm was almost identical to that described in previous studies^13, 23, 50, 55^. The stimulus consisted of drifting bar apertures at various orientations, which exposed a moving checkerboard pattern. The stimulus had a radius of 6.35°, larger than the numerosity mapping stimuli (0.75 radius). Two diagonal red lines, intersecting at the centre of the display, were again presented throughout the entire scanning run. Subjects pressed a button when these lines changed colour, and responded on 80-100% of colour changes within each scanning run.

Visual field mapping data were analysed following a standard population receptive field analysis, as described elsewhere^23, 50^. We identified visual field map borders based on reversals in polar angle of visual field position preference and identified particular visual field maps with reference to previous studies^24, 56–58^.

## Acknowledgements

This work was supported by Netherlands Organization for Scientific Research grants #452.08.008 to SD and #433.09.223 to SD and FW Cornelissen, and by Portuguese Foundation for Science and Technology grant #IF/01405/2014 to BH. The Spinoza Centre is a joint initiative of the University of Amsterdam, Academic Medical Centre, VU University, VU Medical Centre, Netherlands Institute for Neuroscience and the Royal Netherlands Academy of Arts and Sciences.

